# Constant-pH Molecular Dynamics Simulations of Closed and Open States of a Proton-gated Ion Channel

**DOI:** 10.1101/2023.11.30.569372

**Authors:** Anton Jansen, Paul Bauer, Rebecca J. Howard, Berk Hess, Erik Lindahl

## Abstract

Although traditional molecular dynamics simulations successfully capture a variety of different molecular interactions, the protonation states of titratable residues are kept static. A recent *constant-pH* molecular dynamics implementation in the GROMACS package allows pH effects to be captured dynamically, and promises to provide both the accuracy and computational performance required for studying pH-mediated conformational dynamics in large, complex systems containing hundreds of titratable residues. Here, we demonstrate the applicability of this constant-pH implementation by simulating the proton-gated ion channel GLIC at resting and activating pH, starting from closed and open structures. Our simulations identify residues E26 and E35 as especially pH-sensitive and reveal state-dependent p*K*_a_ shifts at multiple residues, as well as side chain and domain rearrangements in line with the early stages of gating. Our results are consistent with several previous experimental findings, demonstrating the applicability of constant-pH simulations to elucidate pH-mediated activation mechanisms in multidomain membrane proteins, likely extensible to other complex systems.

**Significance statement:** Electrostatic interactions play important roles in protein structure and function. Since changes in pH will (de)protonate residues and thereby modify such interactions, pH itself is a critical environmental parameter. However, protonation states of titratable residues are static during classical molecular dynamics simulations. Recently, a *constant-pH* algorithm was implemented in the GROMACS package, allowing pH effects to be captured dynamically. Here, we used this implementation to perform constant-pH simulations of the proton-gated ion channel GLIC, providing insight into its activation mechanism by revealing state-dependent shifts in protonation as well as pH-dependent side chain and domain-level rearrangements. The results show that constant-pH simulations are accurate, efficient, and capable of modeling dozens of titratable sites, with important implications for e.g. drug design.

## Introduction

A prominent problem when studying pH-dependent conformational change is that experimental techniques such as X-ray crystallography and cryo-electron microscopy (cryo-EM) lack the resolution required to determine the protonation states of titratable residues. Even with methods such as neutron diffraction that theoretically can resolve protons [1], the local electrostatic environment in a crystal or on a grid is not guaranteed to correspond to the cellular environment. Given these experimental limitations, computational methods such as molecular dynamics (MD) simulations may provide insight. Although MD is able to capture a variety of both intra- and intermolecular interactions with high precision, traditional simulations typically do not allow for an explicit value of environmental pH. It is generally not possible to model the dynamics of changing protonation states without resorting to quantum mechanics-based approaches that allow for explicit bond formation and breaking, which are prohibitively slow and presently orders of magnitude away from the timescales of large conformational changes in membrane proteins.

Over the years, significant efforts have been invested in capturing pH-dependence in MD simulations in a more realistic fashion [2–29]. To avoid hysteresis effects where the structure adapts to and stabilizes the initially chosen protonation states, some of these *constant-pH* molecular dynamics (CpHMD) methods are based on the concept of λ-dynamics [30] (Fig. 1B). In this approach, a virtual one-dimensional λ-coordinate is introduced for every titratable residue and integrated together with the positions of regular atoms. Each λ-coordinate can move continuously between 0 and 1, with 0 corresponding to the protonated state, and 1 corresponding to the deprotonated state. The charge on the titratable residue is then coupled to the movement of the λ-coordinate, which in turn evolves on a potential specifically chosen as to capture the pH-dependence of the associated residue [28].

**Figure 1:**
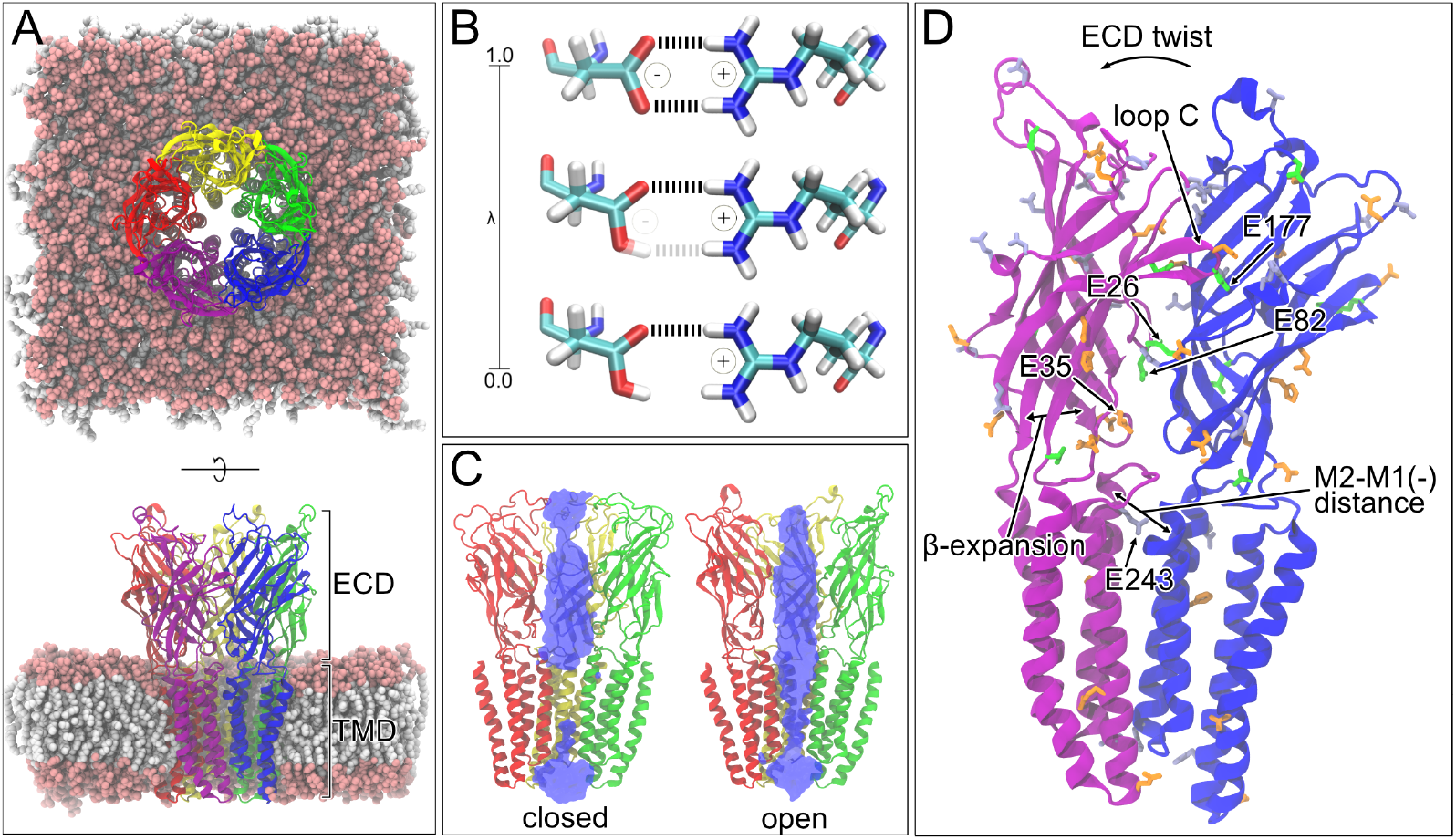
Overview of GLIC structure and CpHMD methodology. **(A)** Top and side view of the GLIC open state embedded in a lipid bilayer, with the ECD and TMD regions indicated. Note the M2 helices lining the central ion-conducting pore. **(B)** Schematic of λ-dynamics based CpHMD for an aspartic acid. Each titratable residue is coupled to a 1D λ-coordinate. As the λ-coordinate changes, so will the charge on the associated residue. **(C)** Closed and open states of GLIC. The front-two subunits have been removed, and pore hydration is shown in blue. In the closed state, it can be seen that the ion-conducting inner pore is dehydrated near the top of the TMD, thereby preventing the passage of ions through the channel. The ions themselves are not shown. During channel opening, conformational changes in the ECD brought on by a lowering of the extracellular pH eventually confer an outward movement of the pore-lining M2 helices, thereby allowing the inner pore to fully hydrate and for the channel to start conducting ions. **(D)** Two opposing subunits in the open state. Titratable residues are indicated, with gray corresponding to residues exhibiting a protonation in accordance with the environmental pH, orange indicating residues that exhibit a perturbed protonation fraction relative to the environment pH, and green for residues exhibiting state-dependent protonation. Loop C and relevant domain-level changes are also indicated, as well as five residues exhibiting major pH and/or state dependence.

Using CpHMD to study large, complex systems such as membrane proteins can be challenging: membrane protein systems frequently contain hundreds of titratable sites, pH-mediated conformational dynamics often involve multiple residues [31], and shifts in protonation tend to be subtle. Furthermore, the protonation states within many groups of such sites are strongly coupled, requiring an integrated approach. However, a new λ-dynamics method recently implemented in GROMACS [32] makes it possible to run constant-pH simulations of an arbitrary number of titratable residues, while still achieving the performance required for sampling at the relevant timescales necessary to explore biological function [28, 29]. This implementation features a fully-consistent particle mesh Ewald (PME) treatment of long-range electrostatics for both the regular atoms and the λ-coordinates [21], a multisite representation for modeling chemically coupled titratable sites such as in histidine (His), and the inclusion of non-interacting buffer particles to maintain a net-neutral system during the simulations. The latter is required to prevent PME artifacts [33].

An interesting family of transmembrane receptors that could benefit from CpHMD simulations are the pentameric ligand-gated ion channels (pLGICs). pLGICs mediate electrochemical signal transduction in a variety of animal cell types [34] and play an especially important role in the nervous system, where they alter the membrane potential upon binding of neurotransmitters. Their involvement in neurotransmission pathways has led to concerted efforts to understand the structure and function of pLGIC sub-families, with several structures now available from X-ray diffraction and cryogenic electron microscopy (cryo-EM) to guide experimental and theoretical studies [35].

One channel that has been of particular interest is the homopentameric bacterial *Gloeobacter violaceus* ligand-gated ion channel (GLIC) (Fig. 1A). Similar to some other bacterial channels where the eukaryotic homologs might be sensitive to voltage or neurotransmitters [36], GLIC is gated by a change in pH, with the channel being closed at neutral pH (7.0) and opening under acidic (pH 4.0) conditions. Despite its name and propensity for modulation by various ligands, GLIC has been primarily shown to be activated upon a lowering of the pH. GLIC is cation-selective, and has served as a model system for the pLGIC family [37], with a multitude of open and closed structures resolved, both in the presence and absence of various ligands and mutations [38–49]. It has also been the subject of multiple computational studies [31, 50–57] to better understand both the gating transition and allosteric modulation. GLIC is an ideal test case, because many, coupled, residues have been experimentally shown to be sensitive to pH. Thus our simulation results can be validated. But details of mechanisms involving several key residues are unkown and therefore constant-pH simulations can provide valuable insights.

Structurally, GLIC assembles as a pentamer that consists of an extracellular domain (ECD) and a transmembrane domain (TMD). The ECD comprises β-strands β1-β10, with loops A-F forming an evolutionarily conserved pocket that functions as the orthosteric binding site in eukaryotic homologs. The TMD comprises α-helices M1-M4, with the M2 helices of each subunit providing the lining for a central ion-conducting pore. Compared to the closed state, the open state of GLIC is characterized by a rigidified and contracted ECD and an expanded transmembrane pore (Fig. 1C). More precisely, the gating transition is believed to proceed from a counter-clockwise rotation of the ECD relative to the TMD (ECD twist), to a contraction between intrasubunit β sheets (β-expansion), and then to a transition of the outward-facing portion of M2 towards the neighboring M1 helix (M2-M1(−) distance) [55]. Among other things, outward dilation of the upper pore-lining M2 helices eventually allows the channel to fully hydrate and start conducting ions (Fig. 1C). Extensive experimental efforts have been invested to identify the site responsible for gating, including full mutational mapping of all Asp, Glu, and His residues in combination with electrophysiology [54], but interestingly it has not been possible to assign the gating process to a single titratable site. Instead, the channel pore opening has been suggested to be driven by an ensemble of protonation events including the pH-sensitive residues E35 and E243 in both experimental and computational assays [31, 54–56] (Fig. 1D). Still, a complete molecular picture of the proton-gating transition of GLIC remains lacking.

In the present work, we demonstrate the applicability of the CpHMD implementation in GROMACS to simulate GLIC in multiple states and conditions, and that it makes it possible to identify sensitive protonation sites and state-dependent charge interactions in the GLIC pH-driven gating process.

## Methods

### System setup

Both closed (PDB ID: 6ZGD [56]) and open (PDB ID: 4HFI [43]) GLIC structures were used as starting models. For 6ZGD, missing side chains were generated using the WHAT IF server [58]. 6ZGD was embedded in a bilayer of 489 1-palmitoyl-2-oleoylphosphatidylcholine (POPC) lipids, while 4HFI was embedded in a bilayer of 498 POPC lipids. Both embedding and subsequent solvation in a 14 × 14 × 15 nm^3^ box were performed using the CHARMM-GUI membrane builder [59]. The CHARMM36-mar2019-cphmd force field [60] was used in combination with the CHARMM TIP3P water model for topology generation. Here, cphmd signifies the inclusion of CpHMD-specific modifications of bonded parameters for titratable residues. Details on these modifications are described in [29]. The phbuilder tool [61] was used to further prepare the CpHMD simulation inputs for GROMACS: all Asp, Glu, and His residues were made titratable. NaCl was added to neutralize the system (at *t* = 0) and ensure an ion concentration of 150 mM, and 185 buffer particles were added to absorb charge fluctuations and maintain a net-neutral system during the simulations (at *t*> 0). For details on the buffer particles, see [28, 62]. Finally, phbuilder was used to generate all CpHMD-specific input parameters.

### CpHMD simulations

All CpHMD simulations were run using the GROMACS CpHMD beta. This version is based on the 2021 release branch and modified to include the routines required for performing the λ-dynamics calculations. The source code branch is maintained at www.gitlab.com/gromacs-constantph until it has been fully integrated into the main distribution.

Energy minimization of the systems was performed using the steepest-descent algorithm. The first relaxation was performed in the NVT ensemble for 10 ps with a time step of 0.2 fs, using the v-rescale thermostat [63] (coupling time 0.5 ps) and a temperature of 300 K. Bond lengths were constrained using the P-LINCS algorithm [64], and electrostatics were performed using the PME method [65]. Subsequent relaxation and production runs were performed in the NPT ensemble using a time step of 2 fs, with pressure kept at 1 bar using the c-rescale [66] barostat (coupling time 5 ps). 30 ns of additional relaxation was performed while gradually releasing restraints on the heavy atoms.

To check whether the default bias potential barriers (7.5 kJ/mol) sufficiently limited sampling of the unphysical λ ≠ 0, 1 states, 20 ns test runs were performed. A number of affected residues were identified, and the inverse Boltzmann method was used to determine the increased barrier height required for those residues. For 4HFI, these included E104 (17.5 kJ/mol), D122 (17.5 kJ/mol), E181 (17.5 kJ/mol), D185 (15 kJ/mol), and E243 (12.5 kJ/mol). For 6ZGD, these included E26 (17.5 kJ/mol), D122 (12.5 kJ/mol), E181 (12.5 kJ/mol), D185 (15 kJ/mol), E243 (12.5 kJ/mol), and E272 (15 kJ/mol).

### Analysis

Mean protonation fractions were obtained by averaging the twenty time-averaged protonation fraction values from the five subunits across the four replicates. Protonation fractions were defined as *N*_proto_*/*(*N*_proto_ + *N*_deproto_), where *N*_proto_ and *N*_deproto_ are the number of simulation frames in which the residue was considered protonated (λ *<* 0.2) or deprotonated (λ *>* 0.8), respectively. A cutoff of 25% was empirically chosen in order to categorize the majority of the titratable residues previously identified as important in terms of their relevance to pH-and/or state-dependent shifts. Simultaneously, this threshold allowed for the exclusion of residues of lesser relevance by assigning them to the unshifted category (Fig. S3). Mean contact occupancies were obtained by averaging the twenty time-averages contact occupancy values from the five subunits across the four replicates. Contact analysis of the acidic residues was performed between either the carboxyl oxygens (Asp, Glu) or the ring nitrogens (His) and the polar (oxygen, nitrogen) atoms of contacting residues. A contact cutoff of 0.40 nm was used for all interactions. In terms of domain-level changes, all measurements were time-averaged over five subunits in four replicas, resulting in bar plots and histograms. The ECD twist was defined as the dihedral angle between the C_*α*_ COM of the ECD (residues 1-192) in subunit *i*, the ECD, the TMD (residues 193 to 315), and the TMD in subunit *i*. β-expansion was defined as the distance between the C_*α*_ COM of residues 30-34 (roughly the β1-β2 loop) and the C_*α*_ COM of residue 190-194 (part of pre-M1 loop) in a single subunit. The M2-M1(−) distance was defined as the distance between the C_*α*_ COM of residues 241-245 (top of M2 helix) in subunit *i* and the C_*α*_ COM of residues 200-204 (top of M1 helix) in subunit *i* + 1. The 9’-radius was defined as the distance between the closest atom in I233 in *a* subunit with the C_*α*_ COM of the I233 residues in *all* subunits. MDAnalysis [67] was used extensively for trajectory analysis, and protein renderings were generated using VMD [68].

## Results

To model state-dependent protonation of GLIC at the atomistic level and demonstrate the applicability of λ-dynamics to this complex pH-gated membrane-protein system, we performed CpHMD simulations of both closed and open structures under resting (pH 7) and activating (pH 4) conditions. As a starting model for the closed state, we used the highest-resolution pH-7 structure available, recently determined by cryo-EM to 4.10 Å (PDB ID 6ZGD) [56]. For the open state, we used one of the highest-resolution pH-4 structures, determined by X-ray diffraction to 2.4 Å (PDB ID 4HFI) [43]. Each subunit of the pentameric channel contains 37 Asp, Glu, and His residues, for a total of 185 per receptor, all of which were made titratable. Each GLIC structure was embedded in a homogeneous lipid bilayer, solubilized in 150 mM NaCl, and set to pH 7 or pH 4. For each of the resulting four systems, four 1-μs trajectories were generated, for a total of 16 1-μs. This required roughly 45 days, using four nodes each containing four Nvidia GeForce RTX 2080 GPUs. We monitored stability particularly in the ECD, where more than 80% of the titratable residues are located; all trajectories converged within 200 ns (Fig. S1). C_*α*_ root-mean-square deviations (RMSDs) of the ECD, excluding flexible loops (residues 1-14 and 49-65), were found to be less than 4.5 Å. As previously observed [56], deviations were particularly low in simulations of the open state (ECD C_*α*_ RMSD < 2.5 Å), likely related to the higher resolution and compacted ECD of this starting structure.

### Constant-pH MD reveals state-dependent shifts in protonation

To quantify pH- and state-dependent protonation shifts in GLIC, mean protonation fractions were computed for all Asp, Glu, and His residues in all four simulation systems (time series provided in Fig. S2). Mean protonation fractions were obtained by averaging the twenty time-averaged protonation fraction values from the five subunits across the four replicates. Protonation fractions were defined as *N*_proto_/(*N*_proto_ + *N*_deproto_), where *N*_proto_ and *N*_deproto_ are the number of simulation frames in which the residue was considered protonated (λ < 0.2) or deprotonated (λ > 0.8), respectively. We then classified each residue relative to the predicted protonation fraction for its amino-acid type in solution, as obtained from potentiometry experiments using alanine pentapeptides [69] and the Henderson-Hasselbalch equation 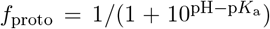 (Fig. 2A). Fifteen acidic residues (D13 on β1, D55 in the β2-β3 loop, E67 and E69 in the β3–β4 loop, D86 and D91 on β5, E104 on β6, D136 in the β7–β8 loop, D145 and E147 on β8, D154 in the β8–β9 loop, E163 on β9, E222 and E243 at the intracellular mouth of the pore, and H277 on the TMD bottom) fell within 25 percentage points of their predicted solution protonation fractions in all four simulation systems (Fig. S3, gray in 1D, Supplemental Table S1). Notably, all of these residues except E243 were located on aqueous surfaces of the receptor, demonstrating how the CpHMD implementation correctly recapitulates the predicted pH-dependent behavior of solvent-exposed regions.

**Figure 2:**
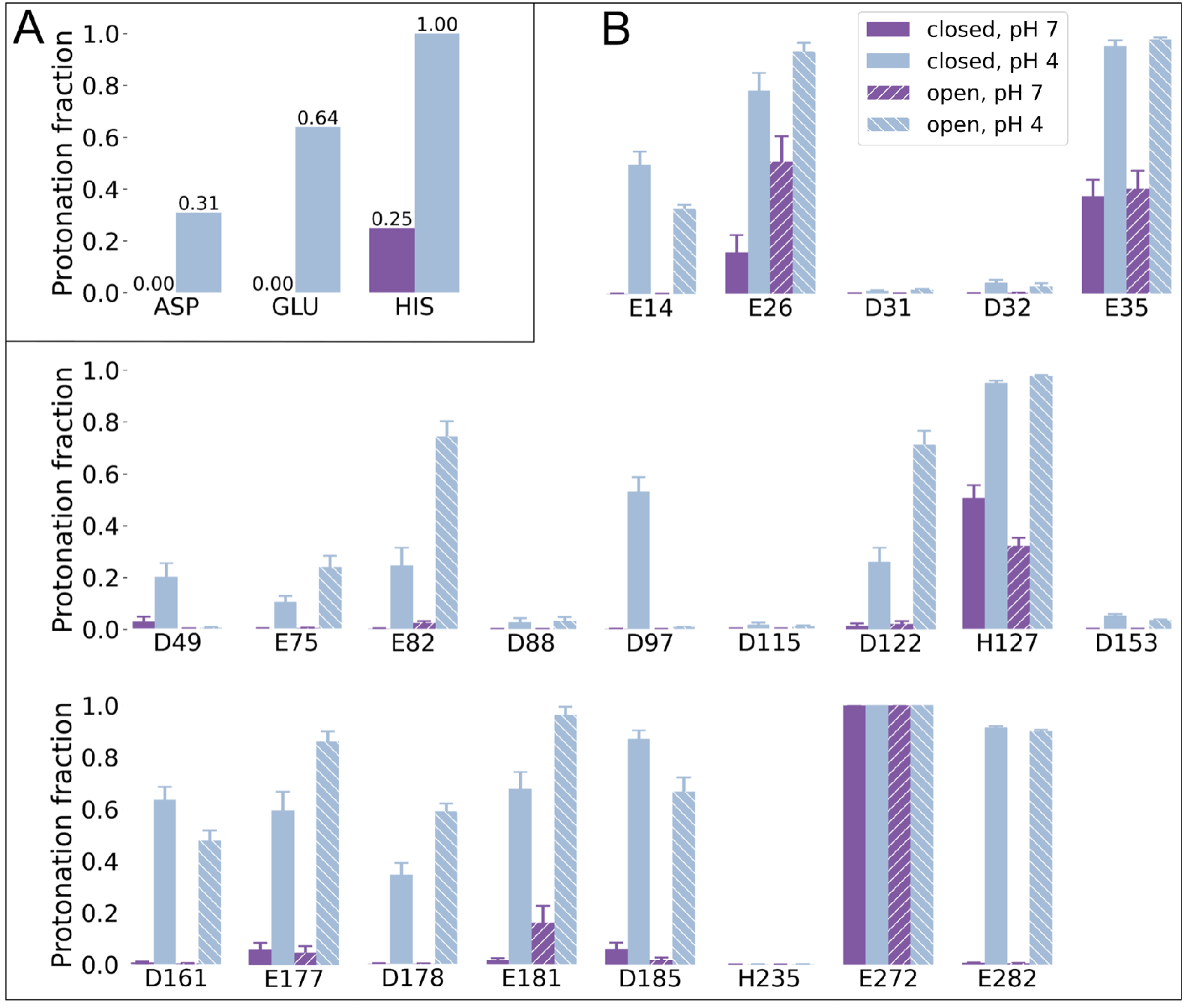
CpHMD reveals state-dependent shifts in protonation. For each titratable residue, mean protonation fractions are provided for both the closed (solid) and open (striped) states at both pH 7.0 (purple) and pH 4.0 (blue), with error bars indicating the standard error estimate. **(A)** Predicted protonation fractions for Asp (p*K*_a_ 3.65), Glu (p*K*_a_ 4.25), and His (p*K*_a_ 6.52) at resting and activating pH, as calculated from potentiometry values for pentapeptides in solution and the Henderson-Hasselbalch equation. This provides a baseline for understanding the shifts observed in panel B. **(B)** Residues exhibiting mean protonations that deviate by more than 25 percentage points from the predicted protonation fraction.

Conversely, 22 buried acidic residues deviated from the theoretical bulk solution protonation by at least 25 percentage points. Of these, several (e.g. E26, E82, E177, E181) also exhibited protonation fractions that differed between the simulated structures, corresponding to either nonconducting or conducting functional states; these are described in detail in the next section. Shifted protonation fractions could be broadly rationalized by contact analysis. Residues with an elevated propensity to protonate (E26 on β1; E35 in the β1–β2 loop; D122 in the β6–β7 loop; D161 on β9; D178, E181, and D185 on β10; E272 and E282 in the TMD) were typically located in the protein interior surrounded by hydrophobic and/or electronegative atoms, environments that favor protonation of acidic residues (Supplemental Table S1). Asp and Glu residues with a low propensity to protonate (E14 on β1, D31 and D32 in the β1–β2 loop, D49 in the β2–β3 loop, E75 and E82 on β4, D88 on β5, D97 in the β5–β6 loop, D115 in the β6–β7 loop, and D153 in the β8–β9 loop) were frequently proximal to one or more basic Arg or Lys residues (Supplemental Table S1), environments predicted to favor their deprotonation.

Despite their complex treatment in CpHMD (due to multiple titratable sites), the three His residues in each subunit converged to distinct protonation profiles, again attributable to local environment (Supplemental Table S1). In the ECD, H127 on β7 was partially buried and exhibited protonation fractions within 26 percentage points of solution predictions (Fig. S3). In contrast, H235 was buried deep within each TMD subunit and did not sample the doubly protonated state (Fig. 2B). Near the intracellular mouth of the pore, H277 was largely solvent-exposed and exhibited protonation fractions within 25 percentage points of solution predictions (Fig. S3).

### Protonation-dependent rearrangements implicated in gating

To refine testable hypotheses for GLIC proton gating based on CpHMD, we visualized the dynamic contacts of several titratable residues in detail (Fig. 3A). We first focused on residues E26 and E35, which exhibited elevated (though submaximal) propensity to protonate even at neutral pH (Fig. 2B). The other highly protonation-sensitive Glu, E272, was more deeply buried between the TMD interior and lipid bilayer, and did not shift protonation states in the experimental range. At E26 on β1, protonation was particularly elevated in open simulations, indicating the protonated form of this residue could selectively stabilize the open state (Fig. 3B). Indeed, although this side chain made relatively few contacts at neutral pH, its protonated form could donate a hydrogen bond to backbone carbonyl atoms of V79 or V81 in the β4–β5 loop of the principal subunit (Fig. 3C, D, E). The resulting intersubunit interaction could promote compacting, twisting motions of the ECD previously implicated in pLGIC gating, as detailed in the next section.

**Figure 3:**
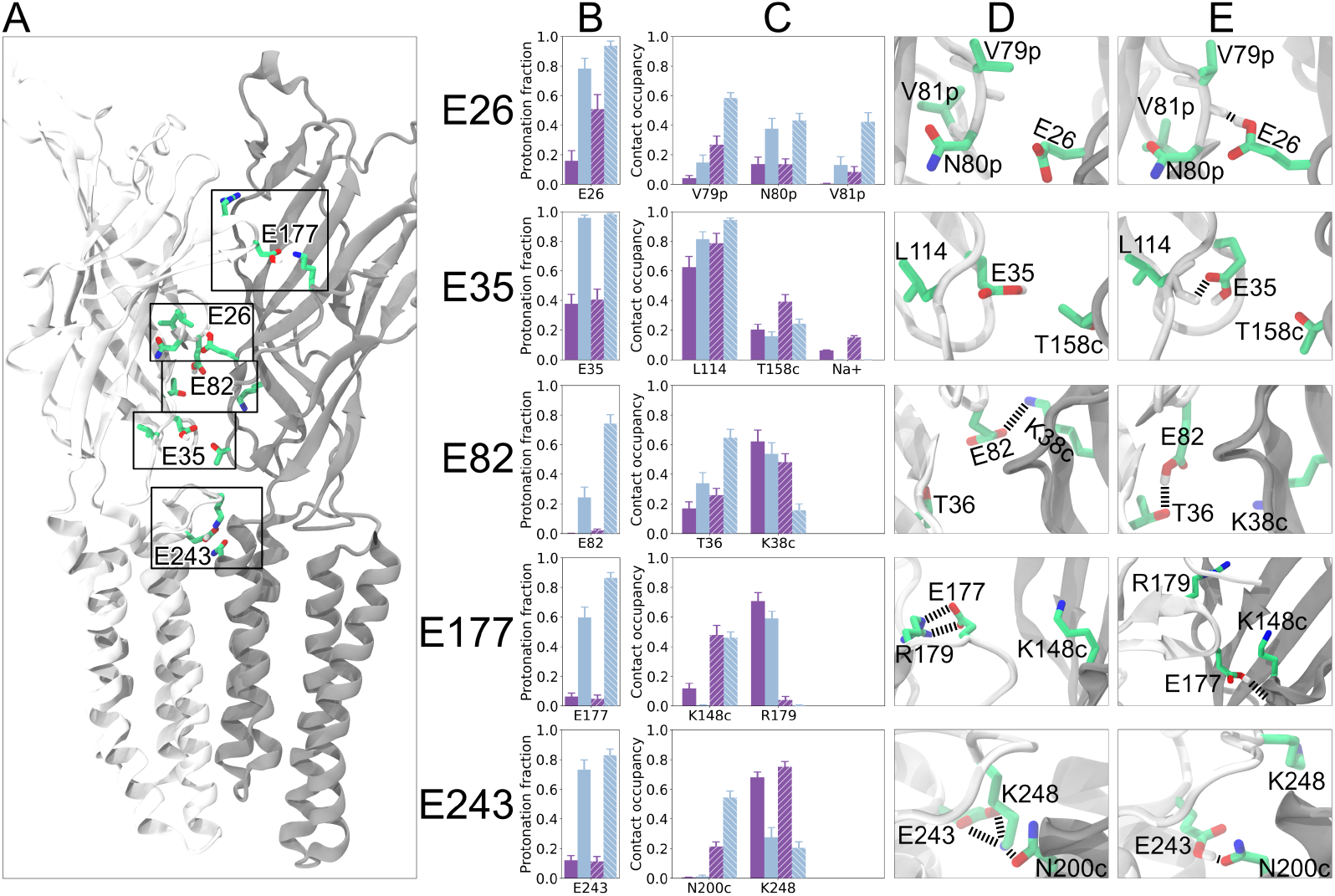
Protonation-dependent rearrangements implicated in gating. This figure shows side chain rearrangements for five residues exhibiting considerable pH and/or state dependence. **(A)** Two opposing subunits shown from the outside of the channel, illustrating the location of residues E26, E35, E82, E177, and E243. **(B, C)** Mean protonation of the titratable residue shown in Fig. 2B **(B)** and hydrogen-bond occupancy of relevant contact partners **(C)**. Data are provided for both the closed (solid) and open (striped) states at both pH 7.0 (purple) and pH 4.0 (blue), with the error bars representing standard error estimates. **(D, E)** Schematics (sticks, colored by heteroatom with dashed black lines corresponding to hydrogen bonds) highlighting spatial differences in contact between the closed state at pH 7.0 **(D)** and the open state at pH 4.0 **(E)**.

Residue E35 exhibited even greater protonation propensity than E26 in the closed state (Fig. 3B), consistent with previous reports that it is an initiating proton sensor [31]. Although state dependence of protonation at this position was limited, polar contacts of E35 with the backbone of L114 in the β6–β7 loop, and with T158 in the β8–β9 loop of the complementary subunit, were modestly increased in the open state (Fig. 3C–E). These interactions would favor contraction of the β1–β2 loop towards β6–β7 and the complementary β8–β9, opposing the β-expansion previously associated with pLGIC closure [55]. These results were also broadly consistent with previous fixed-protonation simulations showing rearrangements of E35 in open versus closed GLIC [56], though under CpHMD, interactions with L114 were favored over those with T158 or sodium ions (Fig. 3C). Sodium ions did bind and unbind frequently within the timescale of all simulations (for example, the mean Na^+^ contact time for E35 in the open state at pH 7.0 was 17 ± 5 ns), providing reassurance that local p*K*_a_ effects of ions are sufficiently sampled.

In contrast to these putative early protonation sites, residue E82 in the β4–β5 loop exhibited relatively little protonation in the closed state, even at pH 4 (Fig. 3B). Indeed, the closed state favors a salt bridge between the deprotonated form of this residue and K38 on β2 of the complementary subunit (Fig. 3C–E). In the open state, rearrangements in this region bring E82 in closer proximity to T36 on β2 of the same subunit, an interaction that favors the protonated rather than deprotonated side chain. Similar to E35, these contacts would oppose β-expansion, instead favoring compaction of β2 towards β4–β5 within a single subunit.

Several acidic residues on β10 (E177, E181, D185) exhibited elevated protonation propensities in the low-pH open state (Fig. 2B). Although this region has not been heavily implicated in GLIC gating, it comprises one strand of Loop C, a critical element that determines neurotransmitter binding to eukaryotic pLGICs [54]. Increased protonation is consistent with closure of β10 over the subunit interface, burying these side chains in a relatively non-polar or electronegative environment in the open state. At the N-terminus of β10, i.e. the tip of Loop C, E177 made a persistent electrostatic contact with R179 on the same strand in the closed state (Fig. 3C–D). In the open state, with the loop relatively closed over the subunit cleft, E177 interacted instead with a backbone carbonyl (of K148) on the complementary β8 strand (Fig. 3C, E), a contact that requires a protonated carboxylic acid side chain. Similar to E26, this rearrangement is expected to promote twisting of the ECD with respect to the TMD.

The M2 residue E243, located at the outer mouth of the transmembrane pore, has also been proposed to influence pH gating. Although protonation propensity did not deviate dramatically from solution values, specific interactions of this residue varied with both pH and functional state (Fig. 3B–C). Consistent with previous reports [56], at neutral pH, E243 was largely involved in a salt bridge with K248 on the M2–M3 loop; this interaction was greatly reduced at low pH, where the side chain was heavily neutralized (Fig. 3B–D). In the open state, E243 interacted with M1 residue N200 from the complementary subunit (Fig. 3C, E), facilitated by the translocation of M2 outward from the channel pore upon activation. The latter interaction was particularly favorable at low pH, where partial protonation of the side chain would facilitate donation as well as accepting of hydrogen bonds.

### Domain-level changes captured by CpHMD

To quantify global as well as local rearrangements captured by CpHMD, we monitored geometric properties of GLIC during simulations of each system (time series provided in Fig. S4, S5, S6, S7). Following conventions of previous computational studies [55, 57], we measured four large-scale changes associated with activation. ECD twist refers to the counter-clockwise rotation of the ECD relative to the TMD and is expected to assume less negative values during activation (Fig. 4A, left). β-expansion refers to the distance between the β1–β2 and pre-M1 loops, and is expected to decrease (Fig. 4B, left). M2–M1(−) distance is measured between the outward-facing ends of the M2 and M1 helices from adjacent subunits, and also decreases with activation (Fig. 4C, left). The 9’ radius indicates the capacity of the permeability filter and is expected to rise (Fig. 4D, left).

**Figure 4:**
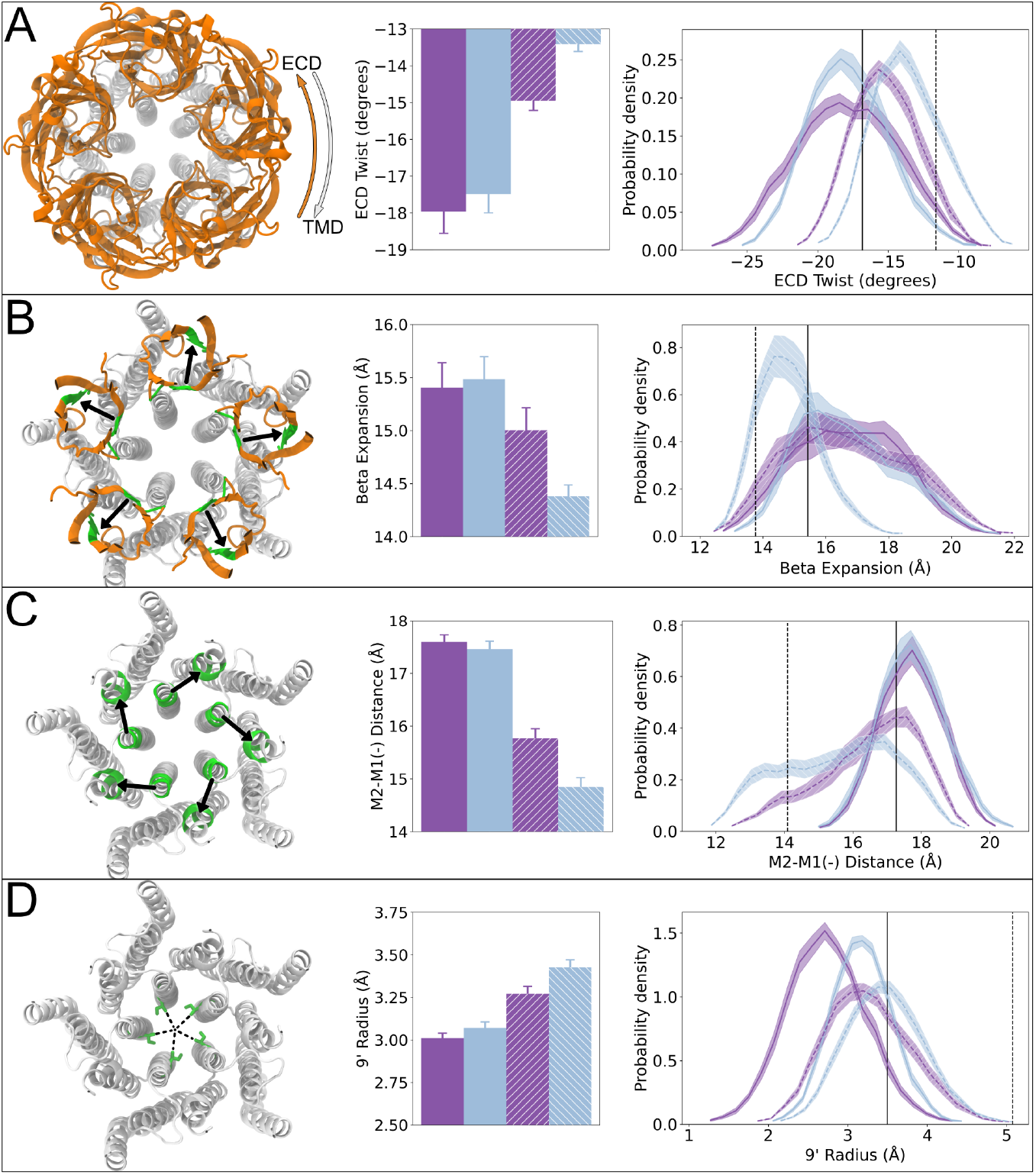
Domain-level changes captured by CpHMD. Four metrics relevant to GLIC gating are shown. For each metric, a motion schematic is provided, with white corresponding to the TMD, orange illustrating the ECD, and green a region of interest. For each metric, mean values and distributions are provided for both the closed (solid) and open (striped) states at both pH 7.0 (purple) and pH 4.0 (blue), with the error bars corresponding to standard error estimates. In the distribution plots, the vertical black solid and dashed lines correspond to the open X-ray (4HFI) and closed cryo-EM (6ZGD) structures, respectively. **(A)** The counter-clockwise twist of ECD relative to the TMD becomes less negative when moving from closed to open states. **(B)** The β-expansion decreases when moving from closed to open states. **(C)** The M2-M1(−) distance decreases from closed to open states. **(D)** The radius at the 9’ position/permeability filter increases from closed to open states.

ECD properties recapitulated experimental data: simulations in the closed state exhibited more negative values of ECD twist and more positive values of β-expansion, with the open state at low pH producing the smallest absolute values (Fig. 4A-B, center). Our simulations also sampled values of both variables corresponding to the initial experimental structures (Fig. 4A-B, right), though the distributions were centered around larger absolute values, indicating a preponderance of closed-like states. The open state at low pH gave a sharper distribution of β-expansion values compared to other systems, indicating a relatively discrete population (Fig. 4B, right). As anticipated, structural transitions between closed and open states were not completed on the simulation timescales; nonetheless, nonequlibrium simulations of the open state raised to neutral pH were associated with increased absolute values of ECD twist and β-expansion, consistent with closing motions.

TMD properties also suggested early stages of gating, indicating that CpHMD simulations are capable of modeling the initiation of the complex process, although simulating complete transitions will (unsurprisingly) require longer sampling. The open state was associated with a relatively contracted M2–M1(−) distance and expanded 9’ radius (Fig. 4, center), although values comparable to the fully open experimental state were not frequently sampled (Fig. 4, right). For M2–M1(−) distance, simulations in the closed state gave sharper distributions, with open-state simulations sampling closed - as well as open-like values. For 9’ radius, all systems shifted towards smaller values than in starting structures, indicating a tendency towards hydrophobic collapse in the channel pore as previously reported [50]. Interestingly, M2-M1(−) expansion and 9’ contraction again shifted more dramatically towards closed-like values in open-state simulations at neutral versus low pH.

## Discussion

Although MD simulations are able to reproduce many types of molecular interactions, they do not typically capture dynamic pH-dependent processes. Instead, the protonation states of titratable residues are usually set at the start of the simulation and remain fixed during the run, no matter the changes in the local environment of such residues. CpHMD methods have made progress addressing this issue, but technical limitations have made them challenging to apply to systems as complex as GLIC due to the large number of titratable, potentially relevant residues involved. The recent GROMACS λ-dynamics CpHMD implementation has made it possible to simulate GLIC under multiple pH conditions including fully dynamic protonation of all 185 Asp, Glu, and His residues, while being fast enough to generate a total of 16 μs of sampling. Despite the system size and number of titratable sites, we previously reported performance was only diminished by approximately 35% compared to classical MD for this system[28], demonstrating the feasibility of the method with contemporary computational resources.

Although complete gating was not observed (nor expected) at these simulation timescales, we were able to capture both pH- and state-dependent shifts in protonation and domain-level rear-rangements in line with predicted early stages of gating. Interestingly, protonation varied occasionally between subunits, but without a clear pattern related to gating and with demonstrably small overall standard errors (Fig. S2). One challenge in our simulations was the tendency of the pore to collapse, limiting our interpretation of transmembrane dynamics and potentially introducing confounding allosteric effects on the ECD. Similar phenomena have been observed in previous MD studies of open ion channels, including GLIC, particularly using CHARMM [50], and might require force field modifications to rectify. Still, the correspondence of several observations here with previous experimental data highlights the applicability of constant-pH simulations for pinpointing early molecular determinants of pH-driven conformational change.

For starting models, we sought structures determined to relatively high resolution with evidence of pH-selective behavior. For the open state, we used an X-ray structure (PDB ID 4HFI) determined at pH 4 to 2.4 Å resolution [43], which has been subjected to classical MD simulations by our group and others under multiple pH conditions [55–57, 70]. Our closed model was a cryo-EM structure determined to 4.1 Å at pH 7 (PDB ID 6ZGD) [56], moderately higher than the best X-ray structure (PDB ID 4NPQ, 4.35 Å) [71], though resolution in cryo-EM and X-ray experiments is not directly comparable. Although an additional cryo-EM structure of the closed state of GLIC was recently reported to an even higher resolution (PDB ID 8ATG, 2.9 Å) [70], this model was derived from cryo-EM data collected under resting, low- and higher-activation pH conditions, rendering the interpretation of pH-sensitive changes uncertain. Regarding the choice of lipid environment, a possible caveat is that POPC does not perfectly represent the native membrane. Since we are however not simulating full gating transitions, and none of the residues discussed in Fig. 3 directly interact with the membrane, we anticipate such differences to have a limited influence on the mechanistic observations in this work.

A sampling difficulty encountered during our CpHMD simulations was optimizing the height of the bias potential for different titratable groups. As described in [28], one of the four potentials applied to the λ-coordinates is a bias potential. The purpose of this is to limit sampling of the unphysical λ ≠ 0, 1 states as much as possible, while still allowing transitions. By default, the height of this potential is 7.5 kJ/mol (see [62] for the effect of the barrier height). In some cases, mainly buried residues engaged in electrostatic contacts, it was observed that these default values were not large enough to prevent sampling of the intermediate λ-states; given the limited maturity of the methods, it is advisable to perform test simulations and adjust barriers before CpHMD production runs. Alternatively, automatic barrier height update schemes could provide a future solution.

Excluding E272, which is buried in the TMD and persistently protonated, E26 and E35 exhibited the highest propensities to protonate at neutral pH out of all Asp and Glu residues (Fig. 2B). Indeed, both E26 and E35 were previously predicted to be sensitive to pH-dependent protonation [55]. ATR/FTIR spectroscopy indicated that E26 changes protonation upon gating [31], and substitutions at this position to Gln or Ala markedly reduced function in electrophysiology experiments [54]. In our simulations, protonation of E26 was associated with increased intersubunit contact with backbone atoms of the principal subunit (Fig. 3), possibly facilitating ECD twisting (Fig. 4A), an early domain-level transition in GLIC gating [55]. E35 has also been implicated as a key proton-sensing residue in GLIC [31], with its mutation dramatically altering function [54]. This residue was indeed sensitive to protonation even at neutral pH; although its protonation state was not demonstrably state-dependent, it may influence gating through state-selective contacts to oppose β-expansion (Fig. 3).

Although our results indicate that E82 and E243 are deprotonated at neutral pH, they both showed state dependence in their protonation and/or contact partners at low pH. E82 is located at the bottom of the vestibular loop and was previously predicted to protonate at low pH [55]. Our data indicate that E82 releases an intersubunit salt bridge in the open state, and instead hydrogen bonds with a neighboring loop in the same subunit, possibly opposing β-expansion to promote activation (Fig. 3). At the extracellular end of the TMD, E243 was previously shown to maintain M2 helix stability [31], while its neighbor K248 was shown to reorient between neutral- and low-pH cryo-EM structures [56]. In our simulations, protonation of E243 at low pH opposes its interaction with K248. No longer locked in a salt bridge, the side chains can reorient, allowing E243 to contact N200 on the complementary subunit (Fig. 3E). This contact could stabilize intersubunit M1–M2 interactions (Fig. 4C) and promote pore opening.

Loop C generally plays an important role in neurotransmitter binding in eukaryotic pLGICs [54]. In GLIC, E177 has been predicted to change protonation states upon gating [31] and to promote opening by opposing ECD spread [55]. In our simulations, Loop-C residues including E177 and E181 were highly protonated at low pH, particularly in the open state (Fig. 2B). E177 could particularly promote opening, as the intersubunit E177-K148 contact was specific to the open state (Fig. 3). Thus, pH sensing in GLIC may involve an evolutionary precursor to classical ligand binding, different in stimulus but mediated by a common structural motif.

## Conclusion

In summary, we have demonstrated the applicability of the GROMACS CpHMD implementation to a complex pH-sensitive membrane protein containing hundreds of titratable residues. Our computational results recapitulate several important experimental findings, including state-dependent protonation of E26 and pH-dependent protonation of E35, as well as side-chain and domain-level rearrangements consistent with GLIC gating. Considering that functional pH sensitivity is exhibited by proteins in the GLIC family and beyond, we anticipate that the present CpHMD implementation will provide a compelling avenue for investigating a variety of complex systems. As the performance overhead of CpHMD is both modest and constant, the time scales such simulations can cover will scale along with improvements in hardware and software, enabling insight into slower processes than are affected by pH.

## Data and code availability

The GROMACS constant-pH beta is available at www.gitlab.com/gromacs-constantph. Trajec-tory subsets, input files, and analysis scripts are archived at https://doi.org/10.5281/zenodo.10869766.

## Author contributions

Conceptualization: AJ, PB, EL, RJH; Methodology: AJ, PB; Investigation: AJ; Data curation: AJ, RJH; Writing - original draft: AJ, RJH; Writing - review and editing: AJ, PB, RJH, BH, EL; Funding acquisition: BH, EL; Supervision: RJH, BH, EL.

## Acknowledgements

This research was supported by the Swedish Research Council (grant nos. 2019-04477, 2019-02433, 2021-05806), the BioExcel Center of Excellence (grant nos. H2020-INFRAEDI-02-2018-823830 and EuroHPC 101093290), the Knut and Alice Wallenberg Foundation, and the Swedish e-Science Research Centre. Computational resources were provided by SNIC (grant nos. 2022/3-40 and 2022/21-16) and NAISS (grant no. 2023/3-27). We thank Pavel Buslaev for constructive support during the project.

