## Supplemental Information for "Constant-pH Molecular Dynamics Simulations of Closed and Open States of a Proton-gated Ion Channel"

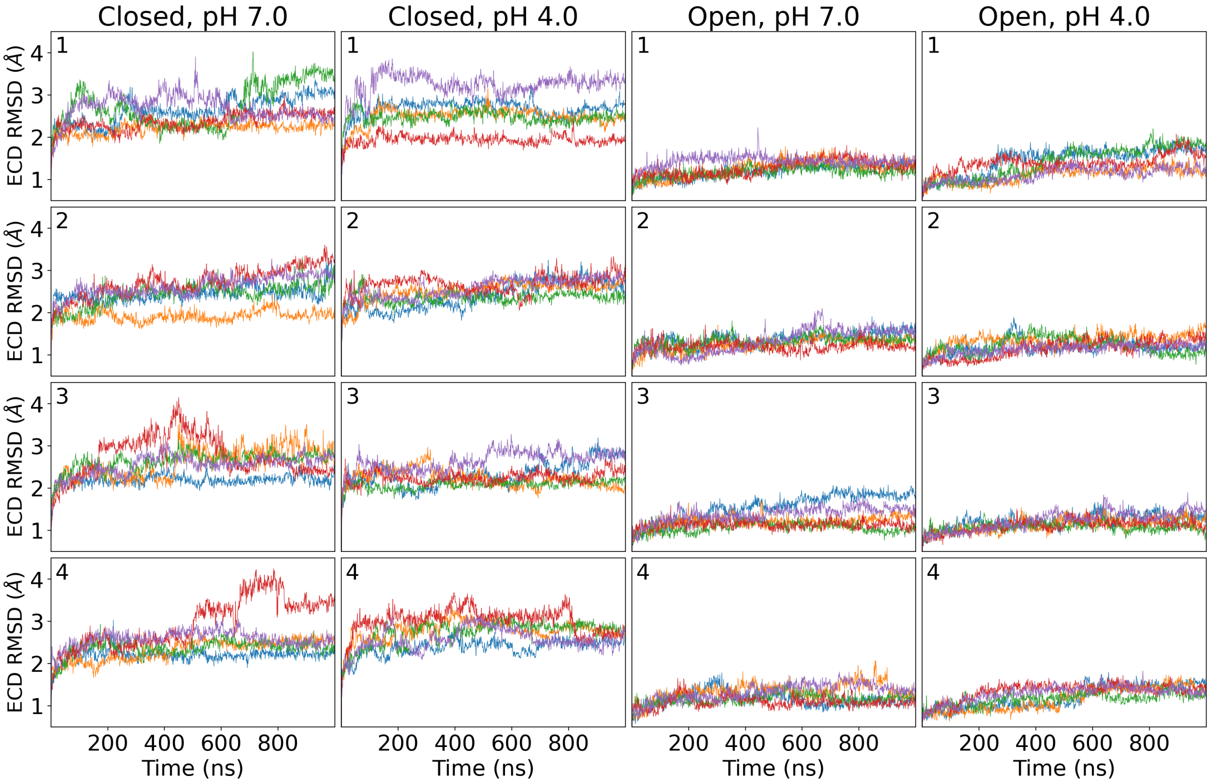


**Figure S1: Root-mean-square deviations for the $C_\alpha$ atoms of ECD residues 15-48, 66-192.** Metric is shown for all simulations (columns), replicates (rows), and subunits (the blue, orange, green, red, and purple curves).

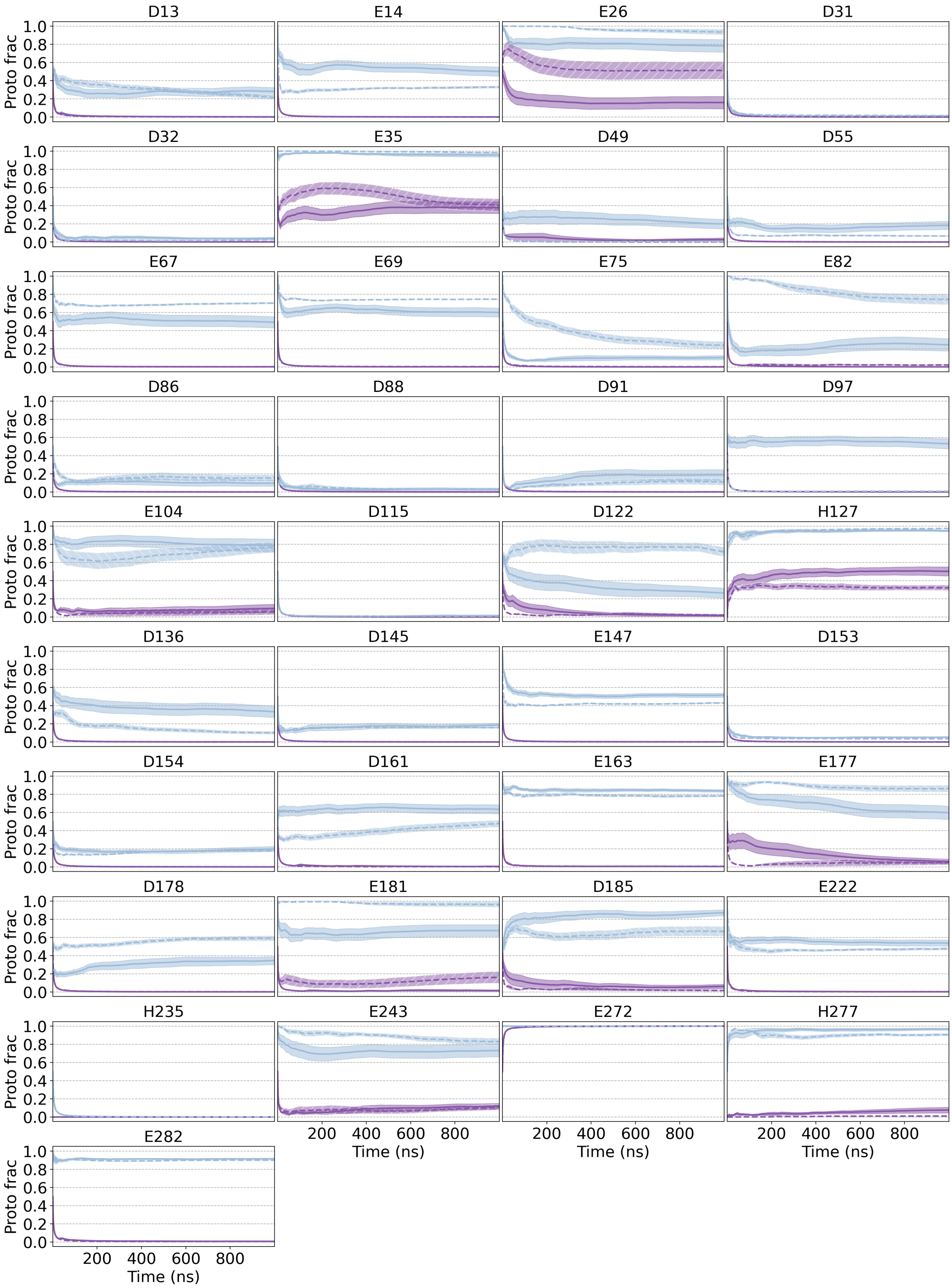


**Figure S2: Time series for the mean protonation fractions of titratable residues.** For each residue, a mean over the cumulative-average protonation fractions of all subunits in all replicates is provided for both the closed (solid) and open (striped) states at both pH 7.0 (purple) and pH 4.0 (blue), with the shaded areas representing the standard error.

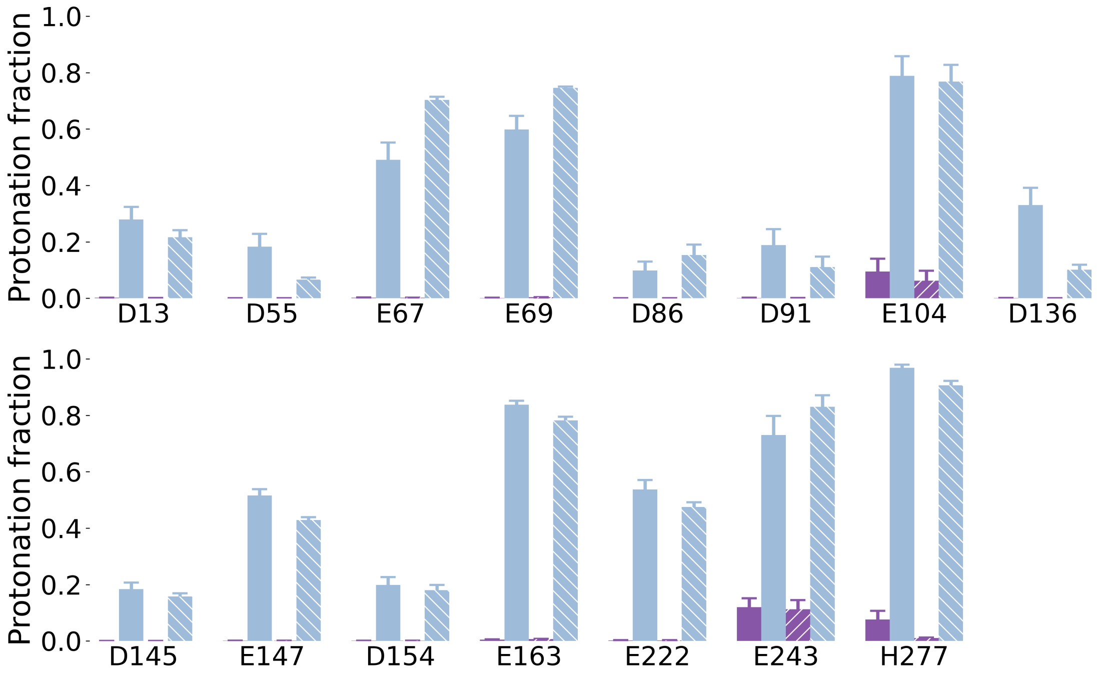


**Figure S3: Residues for which the computed protonation deviates less than 25 percentage points from the protonation fraction associated with the macroscopic pKa.** For each titratable site, mean protonations are provided for both the closed (solid) and open (striped) states at both pH 7.0 (purple) and pH 4.0 (blue), with error bars representing the standard error.

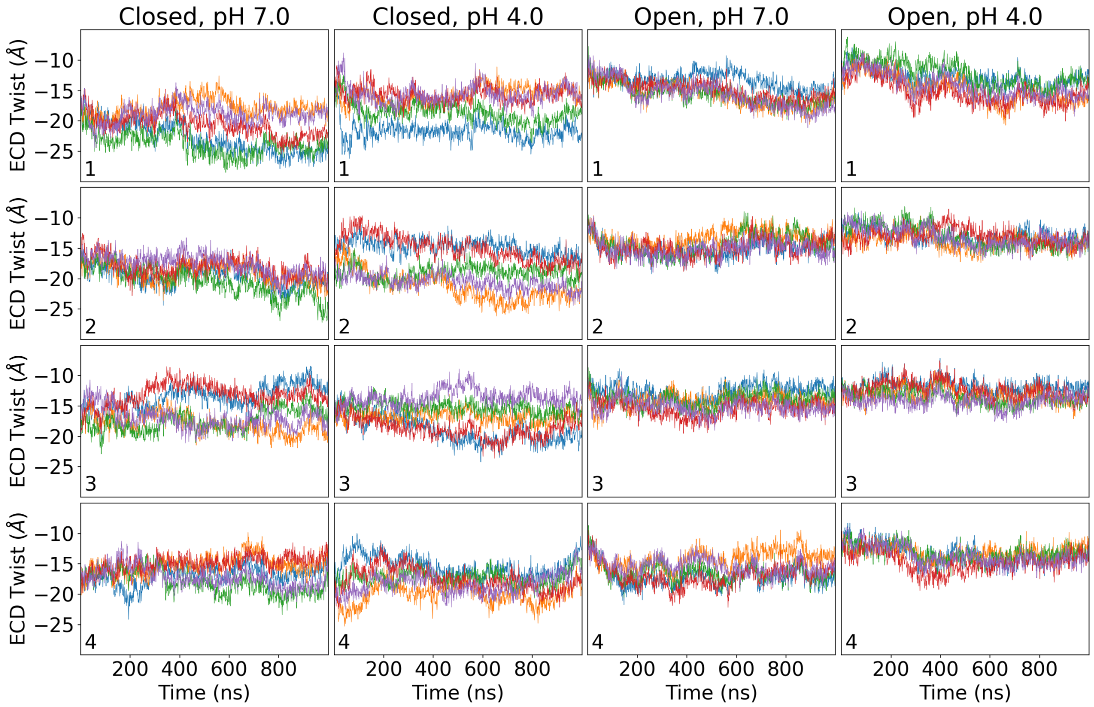


**Figure S4: Time series for the ECD twist.** Metric is shown for all simulations (columns), replicates (rows), and subunits (the blue, orange, green, red, and purple curves).

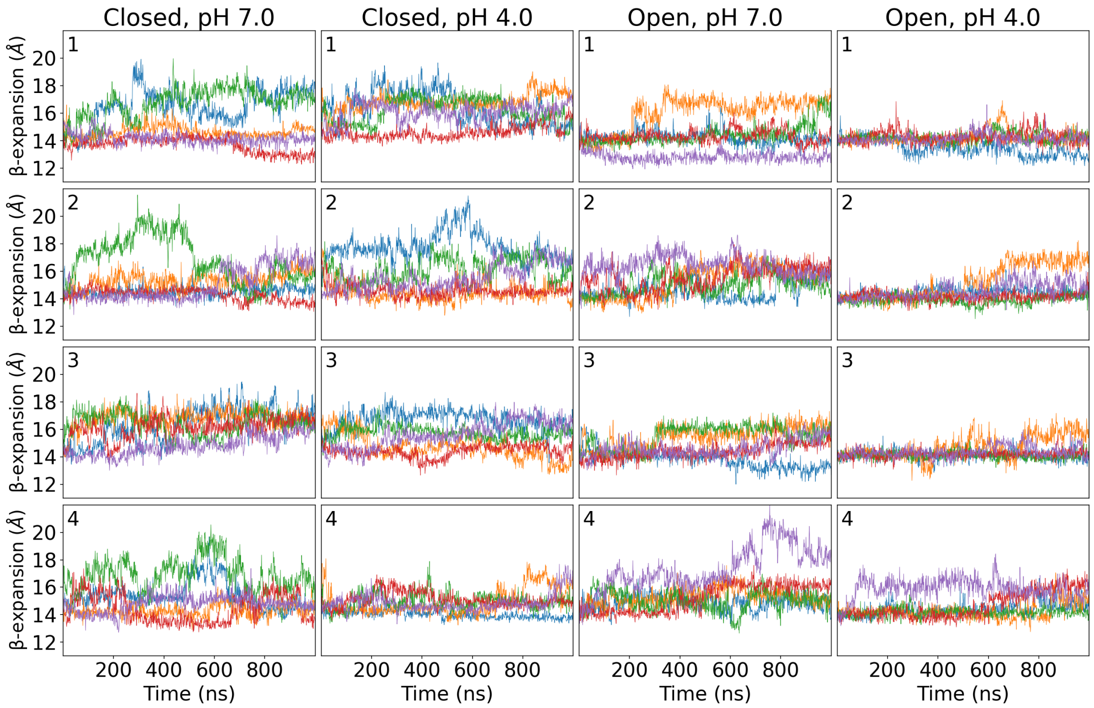


**Figure S5: Time series for the β-expansion.** Metric is shown for all simulations (columns), replicates (rows), and subunits (the blue, orange, green, red, and purple curves).

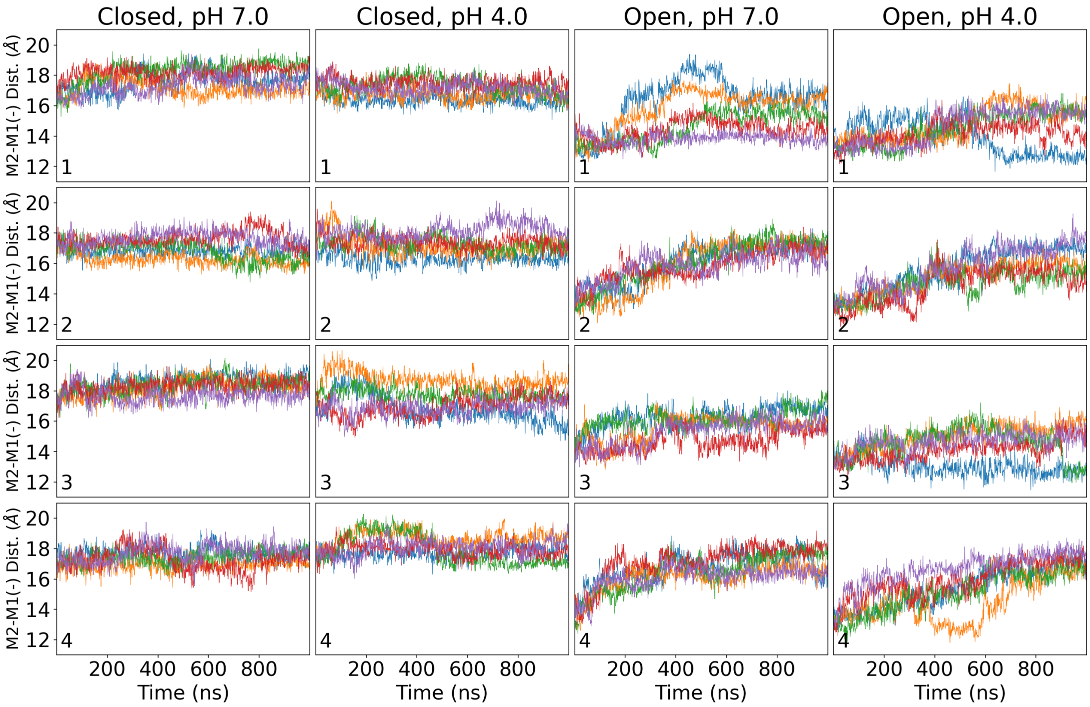


**Figure S6: Time series for the M2-M1(-) distance.** Metric is shown for all simulations (columns), replicates (rows), and subunits (the blue, orange, green, red, and purple curves).

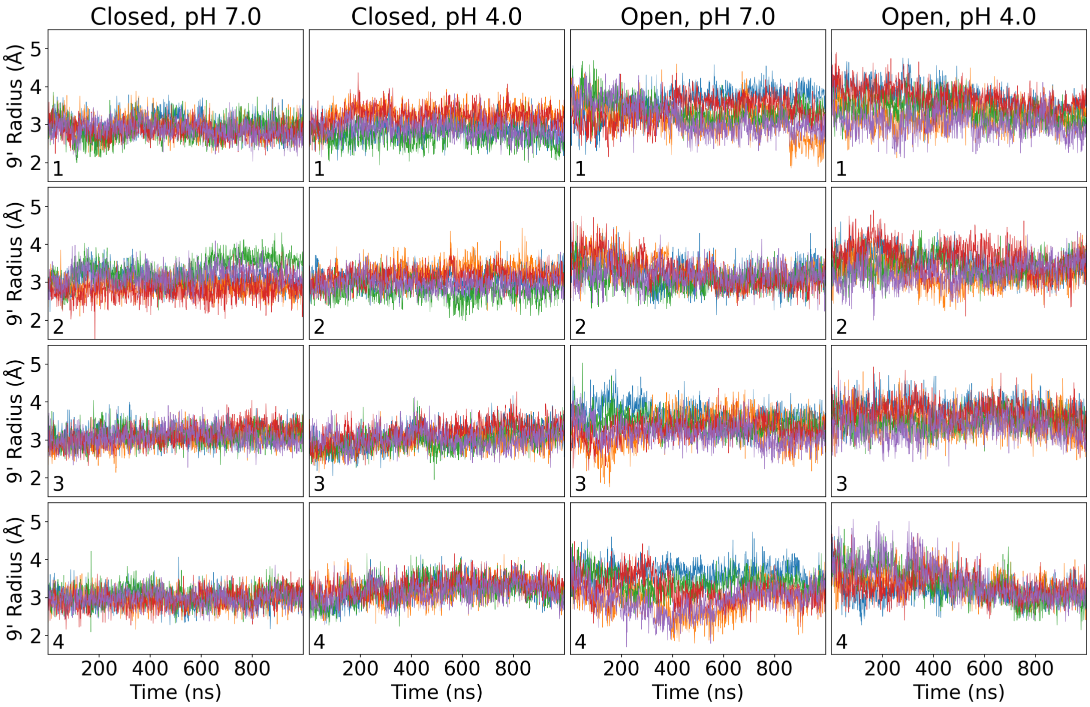


**Figure S7: Time series for 9'-radius.** Metric is shown for all simulations (columns), replicates (rows), and subunits (the blue, orange, green, red, and purple curves).

| Segment | Name | Closed pH 7 | Closed pH 4 | Open pH 7 | Open pH 4 | Δstate pH 7 | Δstate pH 4 | Contact(s) |
|---|---|---|---|---|---|---|---|---|
| $\beta1$ | D13 | 0.00 | -0.03 | 0.00 | -0.09 | 0.00 | 0.06 | Solvent |
| | E14 | 0.00 | -0.14 | 0.00 | -0.31 | 0.00 | 0.17 | R50 |
| | E26 | 0.16 | 0.14 | 0.51 | 0.30 | 0.35 | 0.15 | V79p and V81p in open state |
| $\beta1$–$\beta2$ | D31 | 0.00 | -0.30 | 0.00 | -0.29 | 0.00 | 0.00 | K33 |
| | D32 | 0.00 | -0.27 | 0.00 | -0.28 | 0.00 | 0.01 | R192 |
| | E35 | 0.38 | 0.32 | 0.41 | 0.34 | 0.03 | 0.03 | L114, T158c |
| $\beta2$–$\beta3$ | D49 | 0.03 | -0.11 | 0.00 | -0.31 | 0.03 | 0.20 | R50, R51 |
| | D55 | 0.00 | -0.13 | 0.00 | -0.24 | 0.00 | 0.12 | Solvent |
| $\beta3$–$\beta4$ | E67 | 0.00 | -0.15 | 0.00 | 0.06 | 0.00 | 0.21 | Solvent |
| | E69 | 0.00 | -0.04 | 0.00 | 0.11 | 0.00 | 0.15 | Solvent |
| $\beta4$ (Loop A) | E75 | 0.00 | -0.54 | 0.00 | -0.40 | 0.00 | 0.14 | R77, R133 |
| | E82 | 0.00 | -0.40 | 0.02 | 0.10 | 0.02 | 0.50 | K38c closed, T36 in open state |
| $\beta5$ (Loop E) | D86 | 0.00 | -0.21 | 0.00 | -0.16 | 0.00 | 0.05 | Solvent |
| | D88 | 0.00 | -0.28 | 0.00 | -0.28 | 0.00 | 0.01 | R77p, R105 |
| | D91 | 0.00 | -0.12 | 0.00 | -0.20 | 0.00 | 0.08 | Solvent |
| $\beta5$–$\beta6$ | D97 | 0.00 | 0.22 | 0.00 | -0.30 | 0.00 | 0.52 | S95, G98, T99 |
| $\beta6$ | E104 | 0.10 | 0.15 | 0.06 | 0.13 | 0.03 | 0.02 | Solvent |
| $\beta6$–$\beta7$ (Pro loop) | D115 | 0.00 | -0.29 | 0.00 | -0.30 | 0.00 | 0.01 | R117 |
| | D122 | 0.01 | -0.05 | 0.02 | 0.40 | 0.01 | 0.45 | D115, R118, S123, Q124 |
| $\beta7$ | H127 | 0.26 | -0.05 | 0.07 | -0.03 | 0.19 | 0.03 | Y129 |
| $\beta7$–$\beta8$ (loop B) | D136 | 0.00 | 0.02 | 0.00 | -0.21 | 0.00 | 0.23 | Solvent |
| $\beta8$ | D145 | 0.00 | -0.13 | 0.00 | -0.15 | 0.00 | 0.03 | Solvent |
| | E147 | 0.00 | -0.12 | 0.00 | -0.21 | 0.00 | 0.09 | Solvent |
| $\beta8$–$\beta9$ (Loop F) | D153 | 0.00 | -0.26 | 0.00 | -0.28 | 0.00 | 0.02 | K151 |
| | D154 | 0.00 | -0.11 | 0.00 | -0.13 | 0.00 | 0.02 | Solvent |
| $\beta9$ | D161 | 0.01 | 0.32 | 0.00 | 0.17 | 0.00 | 0.16 | I162, Q193 |
| | E163 | 0.01 | 0.20 | 0.01 | 0.14 | 0.00 | 0.06 | Solvent |
| $\beta10$ (Loop C) | E177 | 0.06 | -0.04 | 0.05 | 0.22 | 0.01 | 0.27 | K148c in open state |
| | D178 | 0.00 | 0.04 | 0.00 | 0.28 | 0.00 | 0.24 | K148c in open state |
| | E181 | 0.02 | 0.04 | 0.16 | 0.32 | 0.15 | 0.29 | V132, R133, R179, L180 |
| | D185 | 0.06 | 0.56 | 0.02 | 0.36 | 0.04 | 0.21 | K183, L184, Y186, Q187 |
| M2 | E222 | 0.00 | -0.10 | 0.00 | -0.17 | 0.00 | 0.06 | Solvent |
| | H235 | -0.25 | -1.00 | -0.25 | -1.00 | 0.00 | 0.00 | N239, I259, Y263 |
| | E243 | 0.12 | 0.09 | 0.11 | 0.19 | 0.01 | 0.10 | K248, N200c in open state |
| M3 | E272 | 1.00 | 0.36 | 1.00 | 0.36 | 0.00 | 0.00 | V273, Q276, S295, T292 |
| | H277 | -0.17 | -0.03 | -0.24 | -0.09 | 0.07 | 0.06 | Solvent |
| | E282 | 0.01 | 0.28 | 0.01 | 0.26 | 0.00 | 0.02 | Y278 |

**Table S1: Protonation fraction analysis data.** On the left, titratable residues and associated segments are specified. Next, the difference between computed protonation fraction and the fraction associated with the macroscopic pKa are provided. Positive deviations $>$ 25 percentage points are colored green while negative deviations $>$ 25 percentage points are colored red. Larger numbers indicate a more perturbed microscopic pKa. Absolute differences in protonation fraction between the closed and open states (Δstate) are also listed. Here, larger numbers indicate more state dependence. Finally, the dominant contact(s) are given.
